# Dynamic mechanisms for membrane skeleton transitions

**DOI:** 10.1101/2024.04.29.591779

**Authors:** M. Bonilla-Quintana, A. Ghisleni, N. Gauthier, P. Rangamani

## Abstract

The plasma membrane and the underlying skeleton form a protective barrier for eukaryotic cells. The molecules forming this complex composite material constantly rearrange under mechanical stress to confer this protective capacity. One of those molecules, spectrin, is ubiquitous in the membrane skeleton and primarily located proximal to the inner leaflet of the plasma membrane and engages in protein-lipid interactions via a set of membrane-anchoring domains. Spectrin is linked by short actin filaments and its conformation varies in different types of cells. In this work, we developed a generalized network model for the membrane skeleton integrated with myosin contractility and membrane mechanics to investigate the response of the spectrin meshwork to mechanical loading. We observed that the force generated by membrane bending is important to maintain a smooth skeletal structure. This suggests that the membrane is not just supported by the skeleton, but has an active contribution to the stability of the cell structure. We found that spectrin and myosin turnover are necessary for the transition between stress and rest states in the skeleton. Our model reveals that the actin-spectrin meshwork dynamics are balanced by the membrane forces with area constraint and volume restriction promoting the stability of the membrane skeleton. Furthermore, we showed that cell attachment to the substrate promotes shape stabilization. Thus, our proposed model gives insight into the shared mechanisms of the membrane skeleton associated with myosin and membrane that can be tested in different types of cells.

**Significance Statement:** Spectrin was first observed in red blood cells, as a result of which, many theoretical models focused on understanding its function in this cell type. However, recently, experiments have shown that spectrin is an important skeletal component for many different cell types and that it can form different configurations with actin. In this work, we proposed a model to study the shared mechanisms behind the function of the actin-spectrin meshwork in different types of cells. We found that membrane dynamics in addition to spectrin and myosin turnover are necessary to achieve conformational changes when stresses are applied and to guarantee shape stability when the stresses are removed. We observed that membrane bending is important to support skeletal structure. Furthermore, our model gives insight into how cell shape is maintained despite constant spectrin turnover and myosin contraction.

## 1 Introduction

To accomplish some of their primary functions, such as motility and cell division, eukaryotic cells need to endure many mechanical challenges (*1*). For example, axons extend long distances and can experience an increase in tension during mechanical deformation. A specific case is the stretch of the sciatic nerves when the ankle flexes due to the specific positioning of the joints (*2*). During normal extension and flexion of the joints, the sciatic nerve has a 5– to 10-fold increase in the strain near the joints (*3*). At the other length scale, red blood cells (RBCs), roughly 8 μm in diameter, deform to go through capillaries and the amount of deformation depends on the shear stress they experience (*4*). The ability of these cells to resist a wide range of deformations is due to the load-bearing features of their structure. Broadly, cell architecture is determined by the canonical cytoskeleton and the membrane skeleton (*5*). The former is a 3D network of filaments, such as actin filaments and microtubules, which provide support to organelles and change their configuration to allow different cell functions. The membrane skeleton consists of a spectrin network beneath the plasma membrane.

Spectrins are proteins that form scaffolds with other molecules inside the cell and confer rapid solid-like shear elasticity to support in-plane shear deformations (*1, 6*). The spectrin scaffold is constructed by attaching the ends of the spectrin rod-like heterotetramers to junctional complexes composed of short F-actin and other proteins (*6*), forming an actin-spectrin meshwork (Fig. 1A). These junctional complexes are one of the structures that connect the spectrin scaffold to the plasma membrane. While cytoskeletal molecules like actin and microtubules use active polymerization to support mechanical loading on cells, spectrin accomplishes its role by either dynamically unfolding or by disassembling the dimer-dimer links (*1, 7, 8*). When a spectrin tetramer is pulled, its repeats unfold and can exhibit a 2.6-fold increase in contour length. The unfolding of the repeats depends on the force and velocity of the pulling (*1, 8*). The structural organization of the spectrin scaffold depends on the cell type (*9*) and as shown more recently, on subcellular location (*10, 11*). In red blood cells, the two main paralogues of spectrin, αI and βI, associate laterally and in an antiparallel manner to form long and flexible heterodimers (*6, 12*). Interactions between the N-terminus of the α-spectrin and the C-terminus of β-spectrin produce bipolar heterotetramers (*6*) (Fig. 1A). The junctional complexes form a pseudo-hexagonal lattice (*6*), which are thought to be regular (Fig. 1B). However, recent experiments showed that F-actin in junctional complexes forms irregular, non-random clusters (*13*).

**Figure 1:**
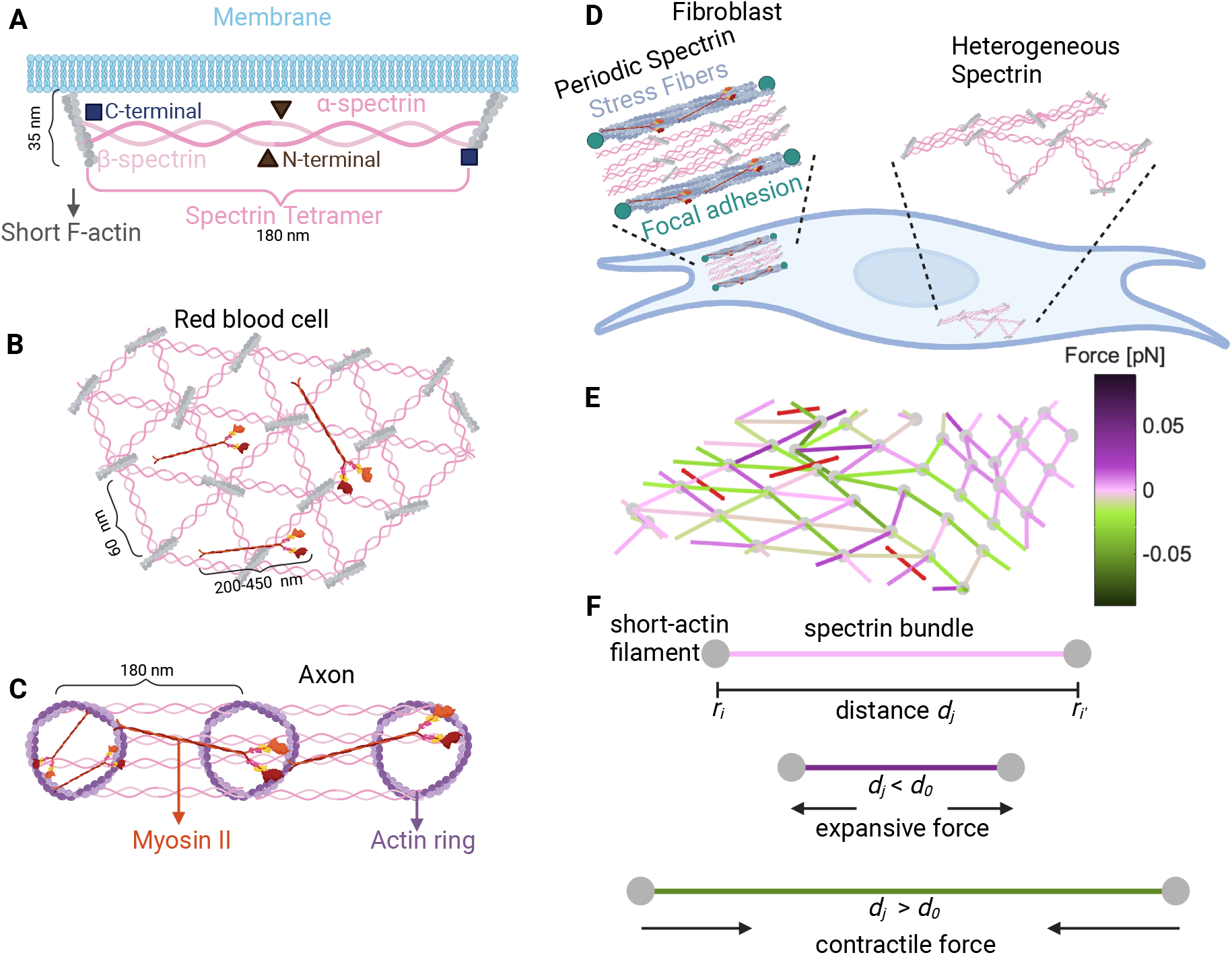
Different configurations of the membrane skeleton. **A)** A spectrin tetramer spanning between short actin filaments. **B**) The hexagonal actin-spectrin meshwork configuration in red blood cells. Myosin generates contractility that may preserve the cell shape (*18*). **C**) Periodic actin-spectrin meshwork configuration in axons. Myosin heavy chains crosslink adjacent actin rings, likely providing tension. Myosin may also span individual rings providing contraction (*15*). **D**) In fibroblasts, the actin-spectrin meshwork has a heterogeneous and dynamic configuration (*11*). **E**) Schematic of the simulated 3D network model. The red lines correspond to myosin, grey nodes to short F-actin, and edges to spectrin, color-coded for the force generated by the spring element. **F**) Schematic representation of the forces generated by the spectrin edges when their length differs from the resting length.

The configuration of the spectrin scaffold in neurons differs in the soma, axon, and dendrite even though it is formed by the same elements, spectrin, actin, and myosin. In axons, α and β spectrin link evenly distributed actin rings, thereby regulating the spacing between rings ( **∼** 180-190 nm) and giving mechanical support to the membrane (*14, 15*) (Fig. 1C). A similar periodic skeleton configuration was found in dendrites (*16*), but the configuration in the soma is similar to that of the RBC (*17*). In fibroblasts (Fig. 1D), spectrin is spatially distributed in regions where the cell edge retracts and there is a low density of actin (*10*). Ghisleni *et al*. showed that the distribution of spectrin is dynamic and it changes during mechanical challenges like cell adhesion, contraction, compression, stretch, and osmolarity changes (*10*). Moreover, recent studies from our group revealed that βII-spectrin transitions between a RBC-like configuration to a periodic axonal configuration in fibroblasts (*11*). Such a transition is driven by actomyosin contractility.

Previously, we used a theoretical model to show that the experimentally observed actin-spectrin transitions in fibroblasts require spectrin detachment from the short F-actin (*11*). Interestingly, some experimental evidence suggests that the actin-spectrin meshwork in the RBC (*7, 13, 18*) and axons (*19*) is also dynamic. In this work, we sought to understand how a minimal system of short actin filaments, myosin motors, and spectrin tetramers can give rise to a wide range of network configurations and confer mechanoprotective capabilities in the cellular context. We used a network model of springs and cables to represent the membrane skeleton (*11, 20–23*) (Fig. 1 E-F) and incorporated the response of the membrane to mechanical stress (*24*). Using this model, we sought to answer the following questions: How does membrane bending interact with the actin-spectrin meshwork? How do myosin contraction and its stochastic addition and removal alter the meshwork? Finally, how do adhered versus detached cells adjust their actin-spectrin meshwork dynamics to conserve their shape? We observed that the balance between the force generated by the bending energy of the membrane and the force generated by spectrin lowers the stress in the membrane. This finding suggests a feedback mechanism between the skeleton and the membrane instead of just the accepted function of the skeleton in providing mechanical support to the membrane. We found that without spectrin unbinding and rebinding to junctional complexes and the action of myosin contraction and its stochastic addition and removal, the actin-spectrin meshwork remains clustered after contractile stress is removed. Therefore, these features of spectrin and myosin are necessary for recovering the pre-stressed configuration of the membrane skeleton. Moreover, our model predicts an optimal number of myosin rods for skeleton recovery from the imposed stress. We showed that, although the membrane skeleton is dynamic, it can maintain cell shape when no stress is induced. We also found that the interplay between the membrane skeleton and the substrate attachments can render stability to adhered cells. We anticipate that our model predictions have implications for a wide-range of mechanoprotective scenarios in which the spectrin-meshwork plays a critical role.

## 2 Results

### 2.1 Qualitative description of the model

We propose a general 3D mesoscopic model for the membrane skeleton to examine its changes in morphology and mechanical properties. This model builts on the 2D model presented in (*11*). The basic component of the model is an actin-spectrin meshwork (Module 1) attached to the extracellular matrix (ECM) through connectors (Module 2), which can induce stress and result in a change in the meshwork configuration. The forces generated by the membrane (Module 3) and myosin (Module 4) also affect the evolution of the meshwork configuration. Thus, a balance between the forces generated by the actin-spectrin meshwork, the membrane, myosin, and connectors dictate the evolution of the meshwork configuration (Module 5). Following (*11*), instead of focusing on exact values for the different model parameters, which are difficult to obtain experimentally and may diverge for different types of cells, we focused on values that allow us to qualitatively represent the meshwork dynamics. Thus, unlike previous modeling efforts that only focus on one type of cell (*24–31*), our model is general. The model parameters are provided in Table 1.

**Table 1:** Model Parameters.

| Symbol | Definition | Units | Value | Reference |
| --- | --- | --- | --- | --- |
| $\zeta$ | drag coefficient | pN s/nm | 1.25 | (11) |
| $\Delta_t$ | time step length | s | 0.002 | (11) |
| **Spectrin** |  |  |  |  |
| $k_{s,S}$ | spring constant | pN/nm | 1 | (11) |
| $d_{0,S}$ | resting length | nm | 180 | (11) |
| $F^{th}$ | force threshold for detachment | pN | 0.05 | (11) |
| **Connecting edges** |  |  |  |  |
| $k_{s,C}$ | spring constant | pN/nm | 1 | fitted |
| $d_{0,C1}$ | resting length for compression | nm | 450 | fitted |
| $d_{0,C2}$ | resting length for extension | nm | 270 | fitted |
| $d_{0,C3}$ | resting length for shear | nm | 270 | fitted |
| $d_{I,C4}$ | resting length for semisphere | nm | 75 | fitted |
| $d_{I,C1}$ | initial length for isotropic stress | nm | 360.1389 | fitted |
| $d_{I,C3}$ | initial length for shear | nm | 1138.4638 | fitted |
| $d_{I,C4}$ | initial length for semisphere | nm | [100,163.05] | fitted |
| **Membrane** |  |  |  |  |
| $k_b$ | bending constant | pN nm | 820<br>($400k_BT$) | (24) |
| $\theta_0$ | spontaneous curvature angle | ° | 0 | fitted |
| $k_A$ | area constant | pN nm | 0.0380<br>($300k_BT/(2d_{0,S}^2)$) | based on (24) |
| **Myosin** |  |  |  |  |
| $k_{c,M}$ | cable constant | pN/nm | 0.1071 | (11) |
| $\varphi_a$ | myosin addition rate | 1/s | 0.01 | (11) |
| $\varphi_r$ | myosin removal rate | 1/s | 0.0063 | (11) |
| $d_{min}$ | minimum length | nm | 135 | (11) |
| $d_{max}$ | maximum length | nm | 450 | (11) |

#### 2.1.1 Module 1: Mechanics of actin-spectrin meshwork

The actin-spectrin meshwork comprises *N*_*e*_ edges connected by *N*_*n*_ nodes, representing spectrin bundles and short F-actin, respectively (Fig. 1E,F). The position of each node, *i*, is given by 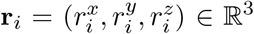, *i* ∈ {1, …, *N*_*n*_}. In what follows, vector quantities are represented using bold letters. The spectrin edges behave like springs with potential *U* ^*spring,S*^, given by

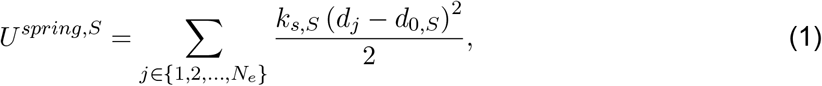

where *d*_0,*S*_ is the resting length and *k*_*s,S*_ is the spring stiffness. The edge *j* spans between the node *i* and *i*^*′*^ and has a length equal to *d*_*j*_ = ||**r**_*i*_ − **r**_*i*′_||. The force generated by *U* ^*spring,S*^ is

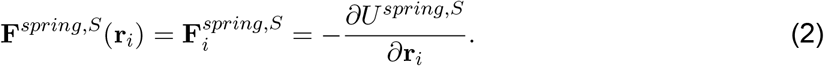

Figure 1F shows the force generated by the spring elements when the length *d*_*j*_ differs from the resting length *d*_0,*S*_: if the edge length is smaller than the resting length, an expansive force is generated, and if the length is larger than the resting length, a contractile force is generated. If the edge length is equal to their resting length, the nodes, which represent actin short filaments, will remain in the same position. See Module 5 for details on the evolution of the position of actin nodes.

### Spectrin unbinding and rebinding

Spectrin dissociates the dimer-dimer links by proteolytic cleavage (*7*). We included the spectrin-spectrin dissociation mechanism in our model by removing the spectrin edges that generate an expanding force greater than a threshold force, i.e.,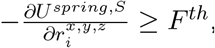, as in (*11*). We also modeled rebinding of the unbound spectrin edges to promote network recovery. Although different rules for spectrin rebinding can be applied, we chose the simplest case, assuming that spectrin tetramers dissociate into dimers at the N-terminal region. Hence, we expect spectrin dimers not to drift away from their current location and be more likely to connect with their previous pair to form tetramers. Moreover, this rule guarantees lower expanding force in the recently connected spectrin edges. Thus, we let the unbound spectrin edges rebind when the distance between the two actin nodes to which an edge was connected equals the resting length *d*_0,*S*_.

#### 2.1.2 Module 2: Induced stress by connection to focal adhesions

To induce stress on the actin-spectrin meshwork, we introduced a new type of spring edge that connects the periphery of the meshwork with fixed nodes representing focal adhesions in the extra-cellular space (Fig. 2A, black lines and circles). These connector edges have spring constant *k*_*s,C*_ and resting length *d*_0,*C*_. We assumed that the focal adhesions are 10 nm lower than the spectrin network in the z-axis, which accounts for the membrane thickness (4-10 nm (*32, 33*)). The initial height difference between the actin and focal adhesion nodes establishes a 3D configuration in the meshwork. The connector edges are attached to spectrin edges through protein complexes, instead of short-actin filaments. Although the protein complexes nodes update their position as described in Module 5, we did not consider these nodes for the membrane forces calculations (Module 3).

**Figure 2:**
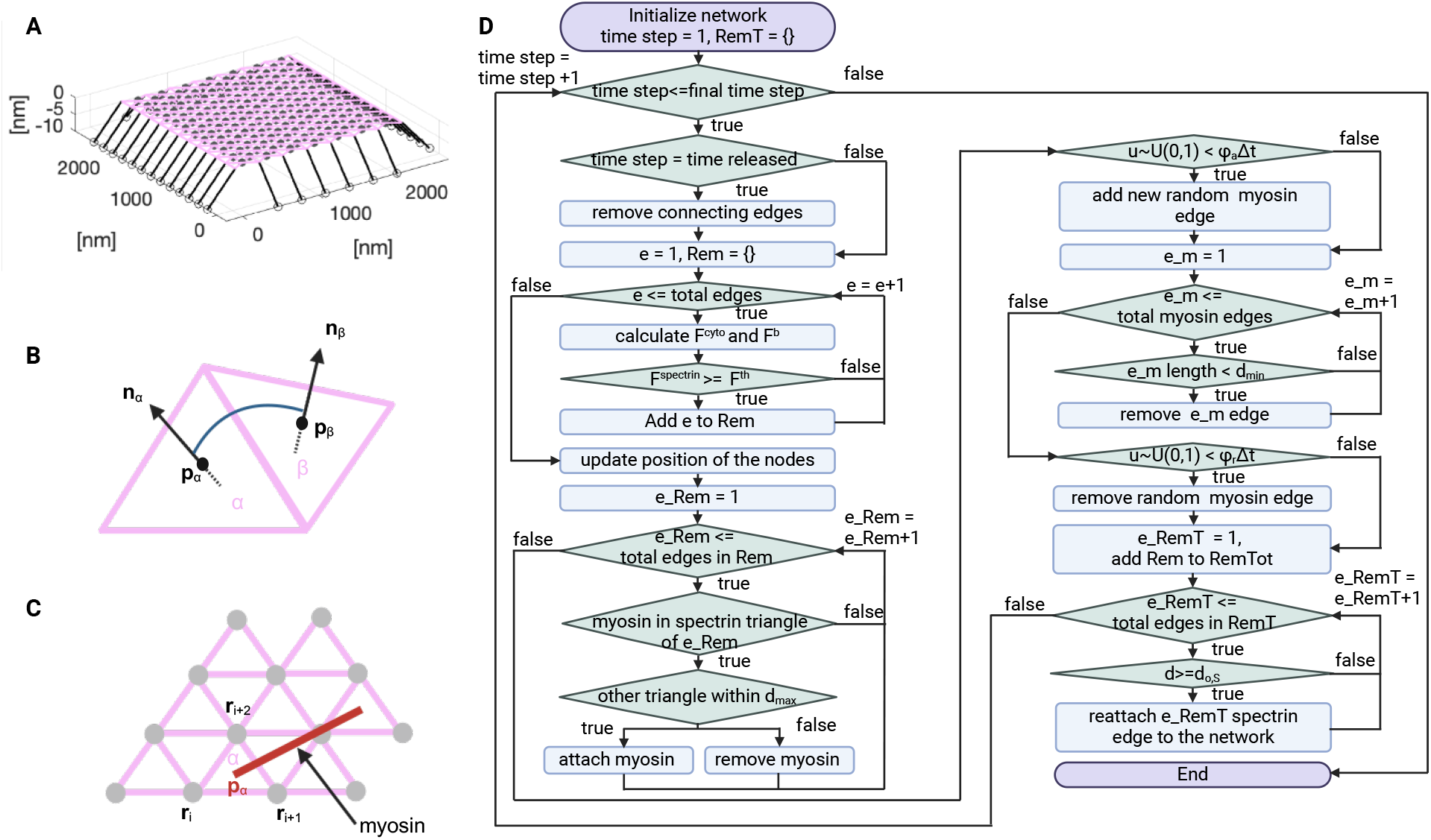
Actin-spectrin meshwork simulation. **A**) 3D view of the initial configuration of the mesh. Pink and black lines correspond to spectrin and connector edges, respectively. Grey filled circles are F-actin nodes and black empty circles represent focal adhesions. **B**) Schematic representation of the angle *θ*_*α,β*_ formed by the *α* and *β* triangular faces of the meshwork. **C**) Myosin edges (red lines) end points are localized at the centers of the spectrin triangles (pink) at position **p**_*α*_ = (**r**_*i*_ + **r**_*i*+1_ + **r**_*i*+2_)*/*3. The force generated by myosin is equally distributed between the F-actin nodes (gray) connecting the spectrin triangle. **D**) Flowchart of the simulation.

#### 2.1.3 Module 3: Membrane forces

##### Bending energy

To model the energy generated by the membrane bending *E*^*b*^, we followed Li *et al*. and assumed that the effects of the lipid bilayer on the cytoskeleton are transmitted via transmembrane proteins and can be represented by coarse-grained local free energies (*24*). Therefore, the bending energy of the membrane affects the short F-actin nodes that are anchored to the membrane. This assumption allows us to use the actin-spectrin meshwork to calculate the bending energy as

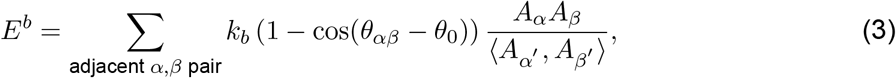

where 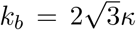, *κ* is the average bending modulus of the lipid membrane (*34*), and *θ*_0_ is the spontaneous curvature angle between two adjacent triangles, *α* and *β*, formed by spectrin bundles. As in (*24*), cos(*θ*_*αβ*_ −*θ*_0_) = cos *θ*_*αβ*_ cos *θ*_0_+sin_*αβ*_ sin_0_, where cos *θ*_*αβ*_ = **n**_*α*_·**n**_*β*_ and sin_*αβ*_ = ±|**n**_*α*_×**n**_*β*_|. Here, sin_*αβ*_ is positive if (**n**_*α*_ − **n**_*β*_) · (**p**_*α*_ − **p**_*β*_)≥ 0. The vectors **n** and **p** represent the normal that points to the exterior and the position of the center of the triangle, respectively (Fig. 2B). Note that in the simulation, the spectrin meshwork can be irregular. Therefore, the contribution of two adjacent triangles is weighted by their area product *A*_*α*_*A*_*β*_ and normalized by the mean product over all the triangle pairs ⟨*A*_*α*′_, *A*_*β*′_⟩ (*24*). Hence, smaller pairs of triangles have less contribution to the bending energy. *E*^*b*^ generates a force 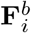, given by,

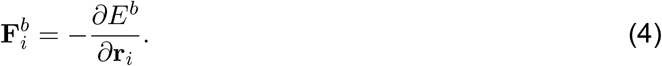

We have neglected the anchorage of spectrin to the plasma membrane through ankyrin for calculating the bending energy. We omitted ankyrin in the model because it binds to the middle of the spectrin tetramer and the model only represents full spectrin tetramers as edges. Thus, considering only short F-actin anchorage at the end of spectrin is sufficient for our simplified representation of spectrin tetramers. Moreover, the function of the spectrin-ankyrin assembly is mostly associated with the organization of membrane proteins in domains (*35*). Hence, we do not expect changes in the membrane bending.

##### Surface area constraint

We assumed that the membrane surface area is conserved in the region of interest and added a surface area constraint. This constraint generates a force

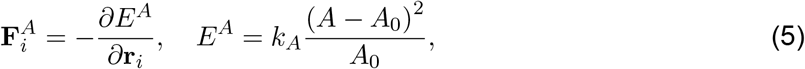

with initial surface area *A*_0_ and area constant *k*_*A*_. Hence, the total force generated by the membrane 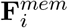 is given by

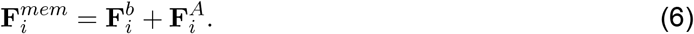

##### Volume exclusion

When simulating closed geometries to mimic cells, we assumed that cells do not shrink indefinitely and implemented volume exclusion in the model that accounts for the organelles and contents of the cytosol. For this, we restrict the movement of F-actin nodes to a volume 15% smaller than the initial volume. If the F-actin node enters this restricted volume, it is reset to its previous value.

Note that our surface area constraint and volume exclusion descriptions do not include the molecular details involved in these processes. We made this simplifying assumption to reduce the complexity of the model. Moreover, mechanisms that regulate membrane surface area and volume have opposite effects (*36*). For example, the membrane surface area regulation by membrane trafficking: On one hand, endocytosis increases the surface area while exocytosis reduces it. On the other hand, an increase (decrease) of membrane tension, which can result from an increase (decrease) of membrane surface area, activates (inhibits) exocytosis and inhibits (activates) endocytosis.

#### 2.1.4 Module 4: Myosin dynamics

We followed (*11*) and added myosin as edges with cable potential energy *U* ^*cable*^, where

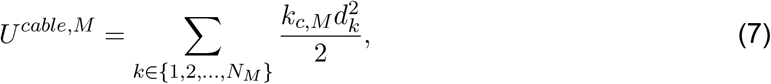

where *k*_*c,M*_ is the tensile force applied by myosin motors and *N*_*M*_ is the number of myosin edges, which are attached to the center of the triangles formed by spectrin edges (Fig. 1C). We assumed that the force generated by the myosin edges 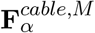 is equally distributed among the three actin nodes joining the triangle formed by the spectrin edges. Thus, the force generated by myosin edges in the F-actin nodes is given by

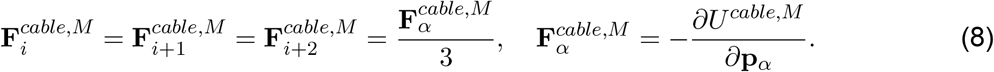

Note that the cable elements only generate a contractile force. If one of the edges of the spectrin triangle is unbound, then the myosin edge tries to attach to a nearby triangle within a distance of *d*_*max*_. If there are none, the myosin edge is removed from the simulation. A myosin edge is also removed from the simulation if its length is less than *d*_*min*_.

##### Stochastic addition and removal of myosin edges

Myosin edges are added and removed from the network randomly at a rate *φ*_*a*_ and *φ*_*r*_, respectively.

#### 2.1.5 Module 5: Evolution of the actin-spectrin meshwork

When stresses are induced to the actin-spectrin meshwork, the actin nodes moves to restore the mechanical equilibrium, given by

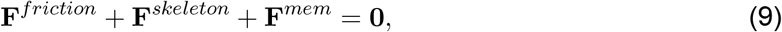

where

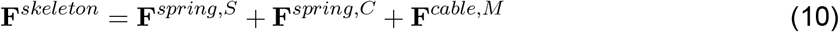

is the force generated by the different elements describing the dynamics of the skeleton and **F**^*mem*^ is the force generated by the membrane (Eq. 6). Note that **F**^*spring,C*^ and **F**^*cable,M*^ act as an external load to the actin-spectrin meshwork, driving it away from equilibrium while **F**^*mem*^ counteracts shape deformations.

Cells are surrounded by other cells and the ECM. Therefore, in Eq. (9), the forces generated by the membrane and skeleton, are balanced by a friction force, **F**^*friction*^. This friction force is created by the viscous dissipation between the movement of the actin nodes and the cell anchorage points to the ECM (focal adhesions) or other cells (cell junctions). Therefore,

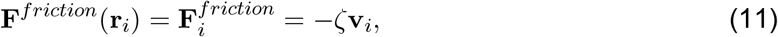

where *ζ* is the drag coefficient and **v**_*i*_ is the velocity at which the actin node at position **r**_*i*_ moves in the absence of friction to restore mechanical equilibrium, i.e.,

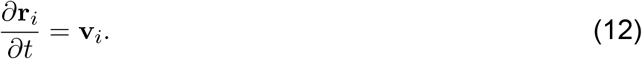

In the simulation, we account for the friction forces (Eqs. 11 and 9). Thus, the evolution of the actin node position is given by

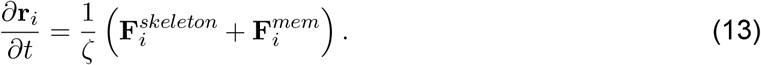

Figure 2D shows the flowchart of the simulation. This simulation framework was implemented in MATLAB and we used it to investigate different scenarios (see Methods for details).

### 2.2 Coupling of membrane bending with the actin-spectrin meshwork is important for resisting isotropic contractility

The membrane skeleton has load-bearing features, which allow cells to resist different deformations. Hence, we tested whether an actin-spectrin meshwork alone can efficiently respond to different imposed stresses. We first simulated the isotropic extension and compression of the actin-spectrin meshwork (Module 1, without spectrin unbinding and rebinding) in the absence of any forces generated by the membrane. We introduced a new type of spring edge (Module 2) that connects the periphery of the actin-spectrin meshwork to fixed nodes representing focal adhesions in the extracellular space (Fig. 2B, black lines and circles). These connector edges are linked to spectrin edges through protein complexes represented by nodes that update their position according to Eq. (13). Therefore, the nodes linking the connector edges with spectrin edges are differentiated from the short-actin nodes (Fig. 3A, black triangles).

**Figure 3:**
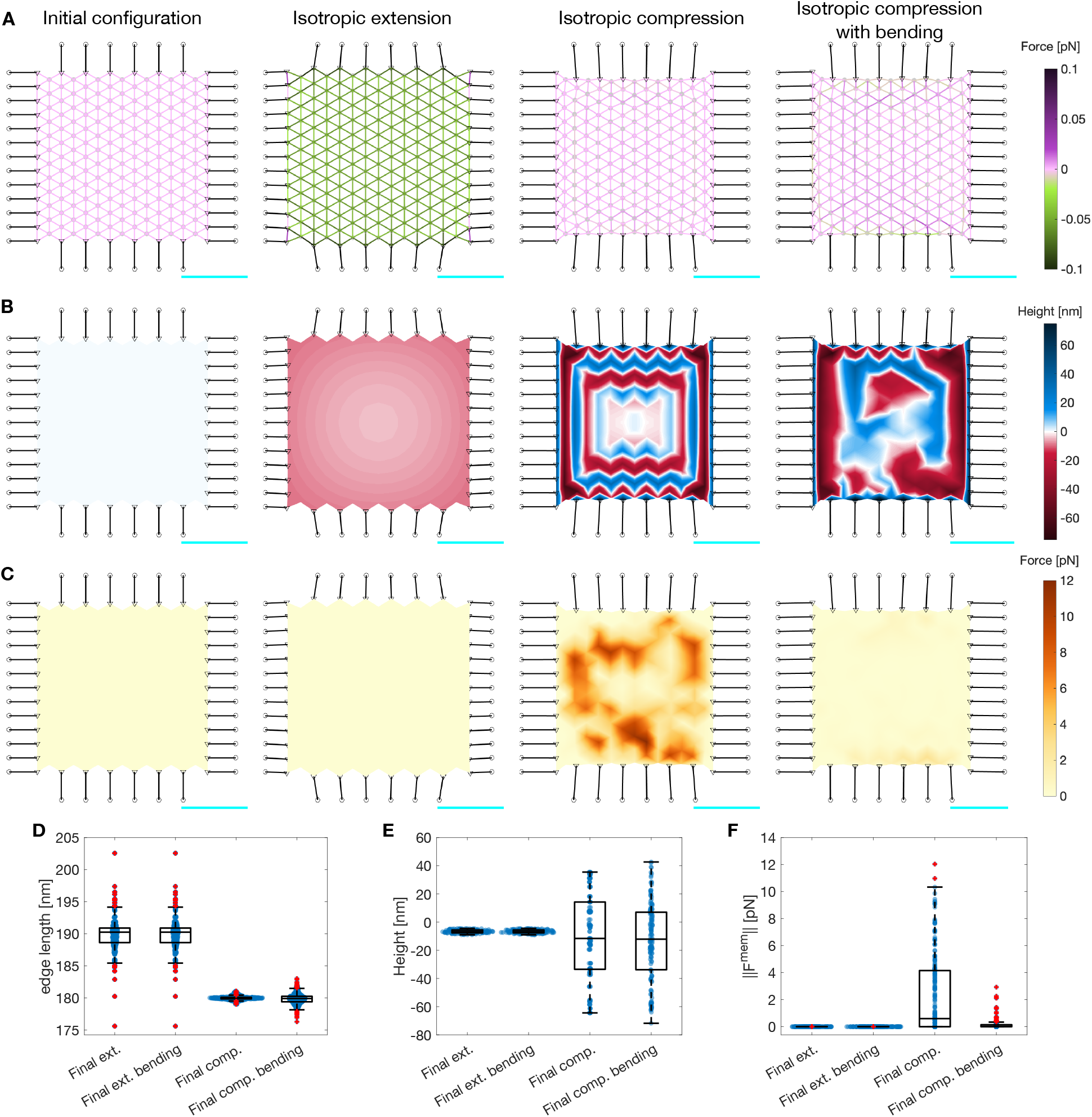
Actin-spectrin meshwork under symmetrical extension and compression. **A**) Configuration of the meshwork initially and after 180 seconds of isotropic extension, compression, and compression including the force generated by membrane bending. The edges corresponding to spectrin edges are color-coded for the force generated by their spring element. The black edges represent connecting edges and the black circles are focal adhesion nodes with –10 nm height. The black triangles are the nodes linking the connector edges to focal adhesions and spectrin edges. The gray nodes show the locations of short F-actin, which have an initial height of 0 nm. The scale bar in cyan corresponds to 1 μm. **B**) Meshwork in A, color-coded for the height of the F-actin nodes. **C**) Meshwork in A, color-coded for the magnitude of the force generated by the membrane 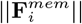. Here 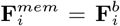, see Methods for simualtion details. Box plot of spectrin edge lengths (**D**), F-actin node height (**E**), and magnitude of the force generated by the membrane (**F**) under the different conditions of (A-C).

We first simulated isotropic expansion. In this case, the connecting edges were pre-extended before the simulation, i.e., we set the initial length, *d*_*I,C*1_, to be larger than the resting length, *d*_0,*C*1_. That way, the connecting edges shrink during the simulation, increasing the contractile force and the edge length of the spectrin edges (Fig. 3A,D). This results in the expansion of the actin-spectrin meshwork. Note that the height of the F-actin nodes decreases at the sides, producing a concave shape of the meshwork (Fig. 3B). To simulate the compression of the meshwork, we set the initial length of the connecting edges smaller than the resting length. We observed shrinkage of the meshwork with a drastic change in the height of the F-actin node locations connecting the spectrin edges (Fig. 3B,E). Note that the length of spectrin edges slightly diverges from *d*_0,*S*_ (Fig. 3D). Therefore, we concluded that the meshwork responded to the isotropic compression by changing the height of F-actin nodes instead of the length of spectrin edges. We determined that the response was induced by the initial difference in height between the actin-spectrin meshwork and the focal adhesion nodes. Moreover, we observed that the magnitude of bending force generated by such height fluctuations (i.e., 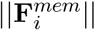, Fig. 3C,F), is high. We concluded that these fluctuations were physiologically unfeasible because they require high amounts of bending energy and the actin-spectrin meshwork by itself was unable to capture isotropic contractility.

We next added the membrane bending force (Module 3) to balance the force generated by the spectrin and connector springs (see Methods). The addition of the bending force to the force balance eliminated the large height fluctuations in the actin-spectrin meshwork (Fig. 3B). We observed that the F-actin node height was closer to the initial height (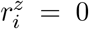 nm) instead of the extreme heights seen in the case without bending (Fig. 3E). As expected, the final configuration minimized the membrane bending energy (Fig. 3C,F). We also found that the bending energy affected the final length distribution of spectrin edges, and therefore, its elastic energy (Fig. 3A,D). However, the addition of the bending energy did not affect the dynamics of the actin-spectrin meshwork under isotropic extension because the difference in actin node height had a slow and smooth evolution (Fig. 3D-F). Thus, our simulations predict that the membrane bending energy interacts with the actin-spectrin meshwork to avoid drastic changes in its configuration when contractile stresses are applied. Furthermore, this interaction minimizes the membrane and spring forces of the actin-spectrin meshwork, resulting in a more efficient physiological response to stresses.

### 2.3 Unbinding of spectrin edges lower stresses due to shear deformation

In cells, spectrin supports in-plane shear deformation (*1*). Hence, we investigated whether our actin-spectrin meshwork with membrane forces can withstand such deformation. To mimic the stress, we removed the horizontal adhesions and changed the position of the vertical adhesions as follows. On the right end of the meshwork, the adhesions were located 360 nm from the linking nodes (black triangles in Figure 4A) in the x-direction and 1080 nm in the y-direction. On the left end of the meshwork, the adhesions were located at similar distances in the x– and y-direction, but with opposite polarity. Since the initial length of the connecting edges *d*_*I,C*3_ is larger than their resting length *d*_0,*C*3_, the connecting edges were pre-extended. This configuration guaranteed that during the simulation, the meshwork would extend in one direction and be compressed in the orthogonal direction (Fig. 4A-C). At the end of the simulation, we observed high fluctuations in short F-actin node height at the center of the meshwork (Fig. 4B), which resulted in high membrane bending force (Fig. 4C). Moreover, the meshwork was under high contractile and expansive forces (Fig. 4A). However, experimental evidence shows that the spectrin network in RBC can deform and experience high shear stress (*7*). To do so, spectrin tetramers dissociate to dimers when a low shearing force is applied (*7*). Therefore, we included this mechanism by removing the spectrin edges that generated an expanding force greater than a threshold force (Module 2), as in (*11*).

**Figure 4:**
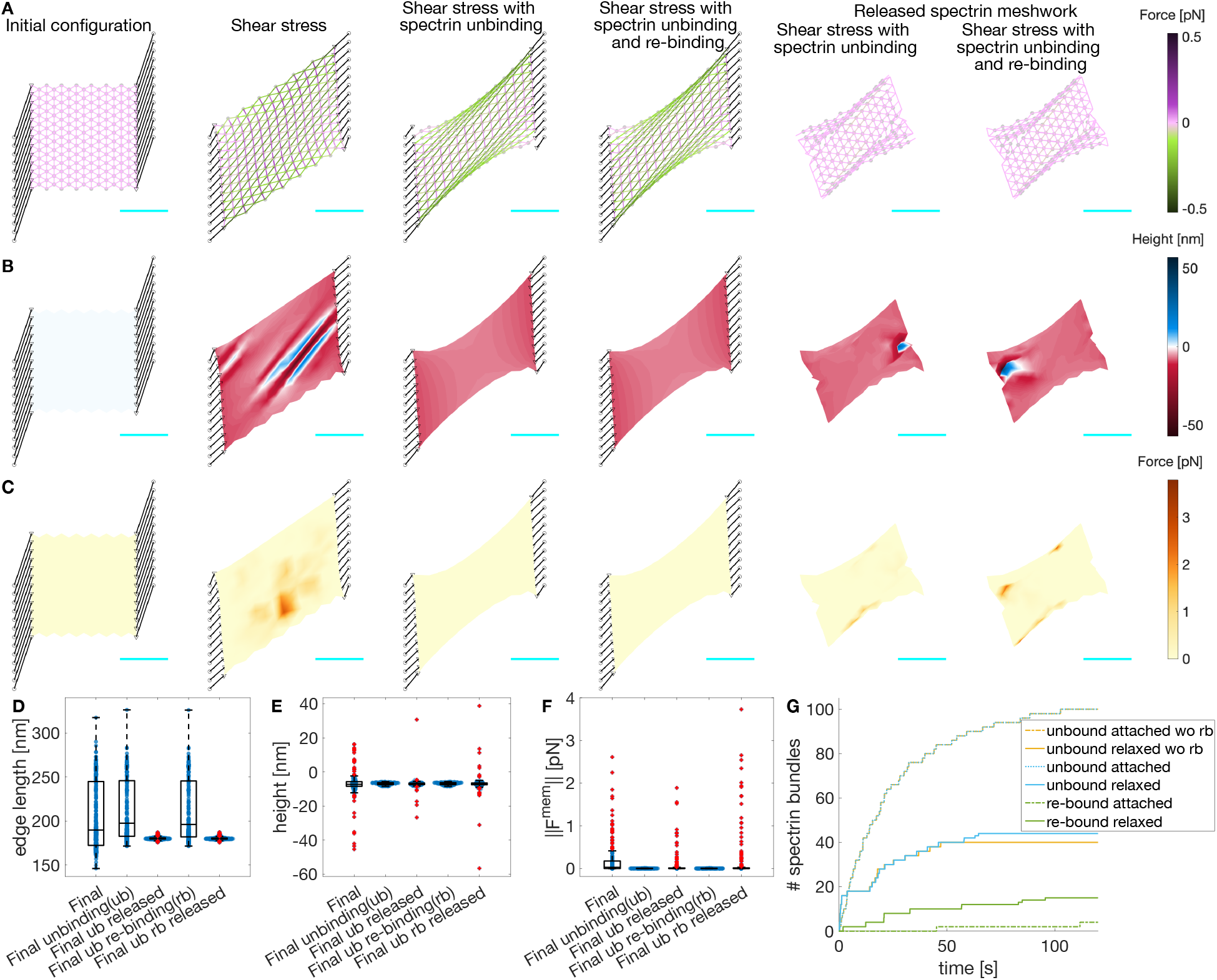
Actin-spectrin meshwork under shear stress. **A**) Initial configuration, and after 120 seconds under shear stress, allowing unbinding of spectrin edges, and allowing unbinding and rebinding of spectrin edges. The last two columns correspond to the case when the meshwork is released from adhesions and evolved for an additional 120 seconds. Edges are color-coded for the force generated by the spectrin spring element. Black lines denote connecting edges and black circles, fixed focal adhesions with –10 nm height. The F-actin nodes (gray dots) and linker nodes (black triangles) have an initial height of 0 nm. The cyan line is a scale bar corresponding to 1μm. **B**) Meshwork on A but color-coded for actin node height. **C**) Meshwork on A but color-coded for the magnitude of the force generated by the membrane. Boxplot of the spectrin edge length (**D**), F-actin node height (**E**), and magnitude of the force generated by the membrane (**F**) distribution under different conditions. **G**) Cumulative sum of the number of unbound and re-bound spectrin edges over time.

Figures 4B,C show that the F-actin height fluctuations and the magnitude of the membrane force were reduced when spectrin unbinding was included in the meshwork dynamics. Furthermore, the final configuration of the meshwork was shrunk along the long axis with large spectrin edges (Fig. 4A). Note that most spectrin removal occurred within the first 30 seconds of the simulation (Fig. 4G, dotted yellow line). At long time, the spectrin edge energy was reduced, reaching a quasi-steady state with little to no spectrin edge removal. Spectrin unbinding reduces mechanical stress, but can the meshwork recover its shape after eliminating the stresses? To test this, we detached the connecting edges from the actin-spectrin meshwork in Figure 4A and simulated for an additional 120 seconds. Note that the meshwork reached a new steady state after 40 seconds with reduced spectrin edge removal (Fig. 4G, yellow solid line). In this new steady state, the resting length of spectrin edges was recovered, thereby minimizing the meshwork stress (Fig. 4A,D). Note that the actin node heights show some fluctuations at the end of the simulation that might result from the force balance (Fig. 4B,E). Such fluctuations also affected the force generated by the membrane (Fig. 4B,F). Overall, the actin-spectrin meshwork remained clustered on the long axis sides and did not recover its shape.

We next investigated whether spectrin rebinding would change the response of the meshwork. We allowed the unbound spectrin edges to rebind when the distance between the two F-actin nodes to which an edge was connected was equal to the resting length. Figure 4A-C shows that the evolution of the meshwork with spectrin unbinding and rebinding is similar to that of the meshwork with only unbinding. Moreover, the evolution of the total number of spectrin unbound edges was similar when the meshworks were attached to the focal adhesions (Fig. 4G dotted yellow and blue lines). The resulting meshworks only differed when released from the focal adhesions: the meshwork allowing spectrin edge rebinding showed more unbinding events (Fig. 4G solid yellow and blue lines). As expected, spectrin rebinding events were fewer when the meshwork was attached to focal adhesions, but the events increased when it was released (Fig. 4G green). However, when the meshwork reached a steady state, i.e., the length of spectrin edges was equal to the resting length, the unbinding and rebinding events ceased and the meshwork remained clustered. Hence, we hypothesized that additional mechanisms are needed to prevent spectrin clustering after the stress is removed, thereby promoting spectrin redistribution in the cell to provide mechanical support to the membrane and bear future stresses at different locations.

### 2.4 Myosin interactions with the actin-spectrin meshwork promote recovery of spectrin edges

We next asked under what conditions would the membrane skeleton recover a prestressed configuration after the external loading is removed. Recent work has shown that spectrin topological transitions are driven by actomyosin contractility (*11*). Therefore, we incorporated the dynamics of myosin into the meshwork (Module 4). As in (*11*), myosin edges generated a contractile force and were removed when they shrunk to a minimal length or when there were no available binding sites. In addition to these dynamics, myosin edges were added and removed stochastically.

We observed that the length of spectrin edges and height of actin nodes were similar to that of the meshwork without myosin (Fig. 5A-F). However, myosin increased the rebinding events after the meshwork was released (Fig. 5G). Due to the stochastic nature of the myosin dynamics, we ran 30 additional simulations to test the generality of the results and obtained statistics. We observed that after the initial 60 seconds, the number of myosin edges in the system with a surface area of ≈ 2.5 μm^2^ settled to one (Fig. 5H), which promoted unbinding and rebinding in the spectrin meshwork (Fig. 5G). Thus, a single myosin edge per 2.5 μm^2^ was enough to promote spectrin edge turnover. In some simulations, the final percentage of attached spectrin edges matched the percentage before releasing the spectrin meshwork from focal adhesions (Fig. 5I). We concluded that myosin addition avoids the clustered, crumpled state and helps meshwork recovery, which prepares the membrane skeleton to respond to new stresses.

**Figure 5:**
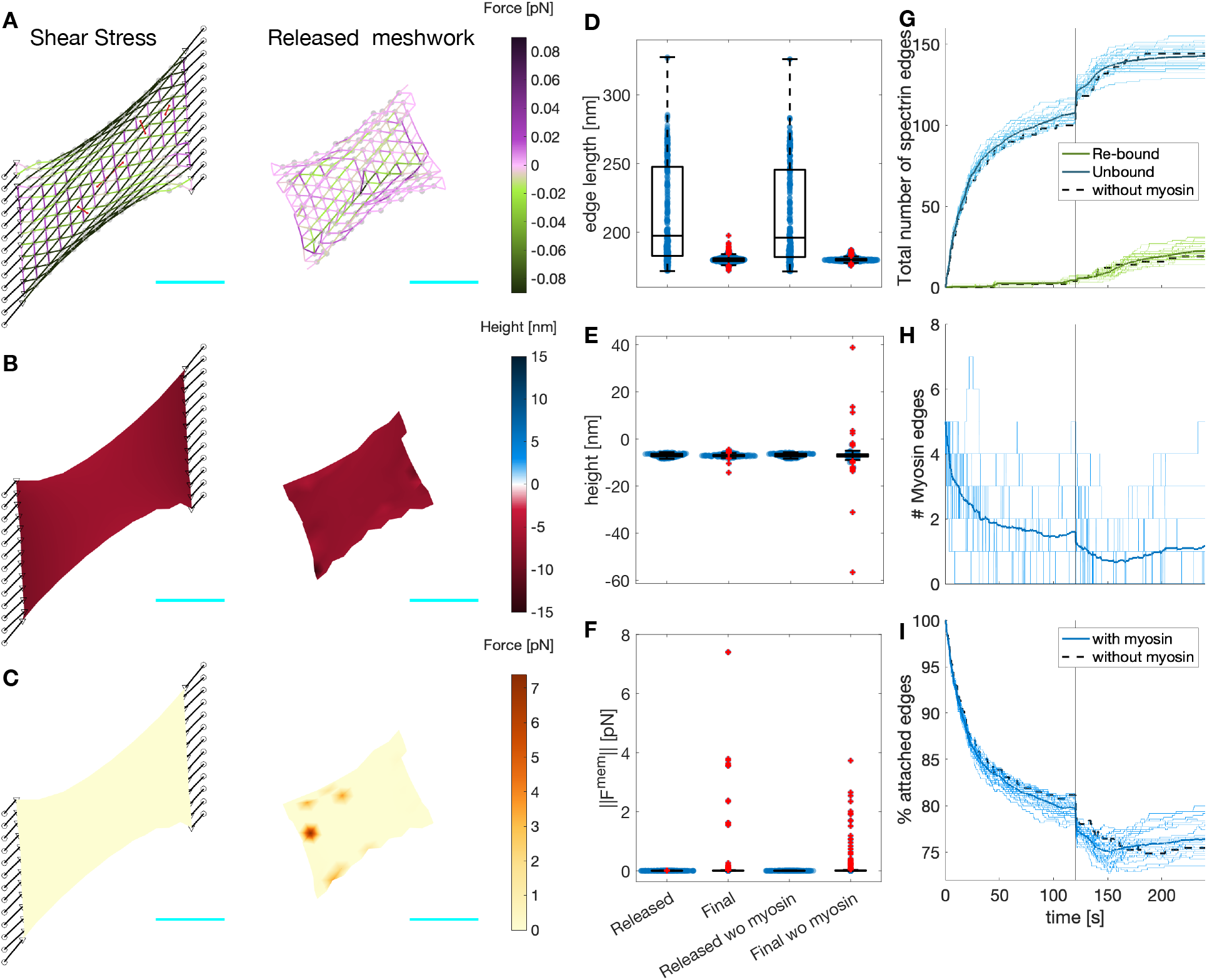
Myosin dynamics on an actin-spectrin meshwork under shear stress. **A**) Meshwork configuration after 120 seconds under shear stress, and 120 seconds after releasing the network from the focal adhesion nodes. Edges are color-coded for the force generated by the spectrin spring element. Black lines denote the connecting edges and black circles fixed focal adhesions with –10 nm height. Red edges correspond to myosin. Gray circles represent F-actin nodes and black triangles, linker nodes with an initial height is 0 nm. The cyan line is a scale bar corresponding to 1μm. **B**) Meshwork on A but color-coded for actin node height. **C**) Meshwork on A but color-coded for the magnitude of the force generated by the membrane. Boxplot of the spectrin edge length (**D**), actin node height (**E**), and membrane force magnitude (**F**) for the configurations in A-C. The values for the meshwork without myosin (Fig. 4) are given for comparison. **G**) Evolution of the total number of unbound (blue) and re-bound (green) spectrin edges. **H**) Evolution of the number of myosin edges. **I**) Evolution of the total number of attached spectrin edges. In G-I, the thin lines correspond to 31 different simulations, the thick line is the temporal average of the simulations, and the black dotted line shows the evolution of the meshwork without myosin in Figure 4.

Next, we investigated whether increasing the number of myosin edges acting in the actin-spectrin meshwork enhanced the rebinding of spectrin edges, thereby, the meshwork recovery after removing the stress. For this, we changed the ratio of rates corresponding to the random addition and removal of the myosin edges. This rate ratio, (*rr*), given by

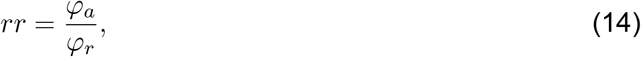

where *φ*_*a*_ and *φ*_*r*_ are the rates for random addition and removal of myosin edges, respectively. We found that increasing (decreasing) the rate ratio results in more (less) myosin edges in the simulations, even after releasing the meshwork from the focal adhesions (Fig. 6A). Moreover, increasing *rr* raised the median and reduced the spread of the myosin edges lifetime (Fig. 6D). Thus, the higher the stochastic addition-to-removal ratio, the more myosin edges exert contraction at different zones of the actin-spectrin meshwork. Paradoxically, this hinders the contraction of the myosin edges and their removal when they reach a minimal length, extending the edge lifetime but preventing the creation of space for spectrin edges to rebind. On average, both increasing and reducing *rr* resulted in a smaller increase in the percentage of attached spectrin edges (Fig. 6B,C), suggesting that there is an optimum number of myosin edges acting on the meshwork that allows further rebinding of spectrin edges.

**Figure 6:**
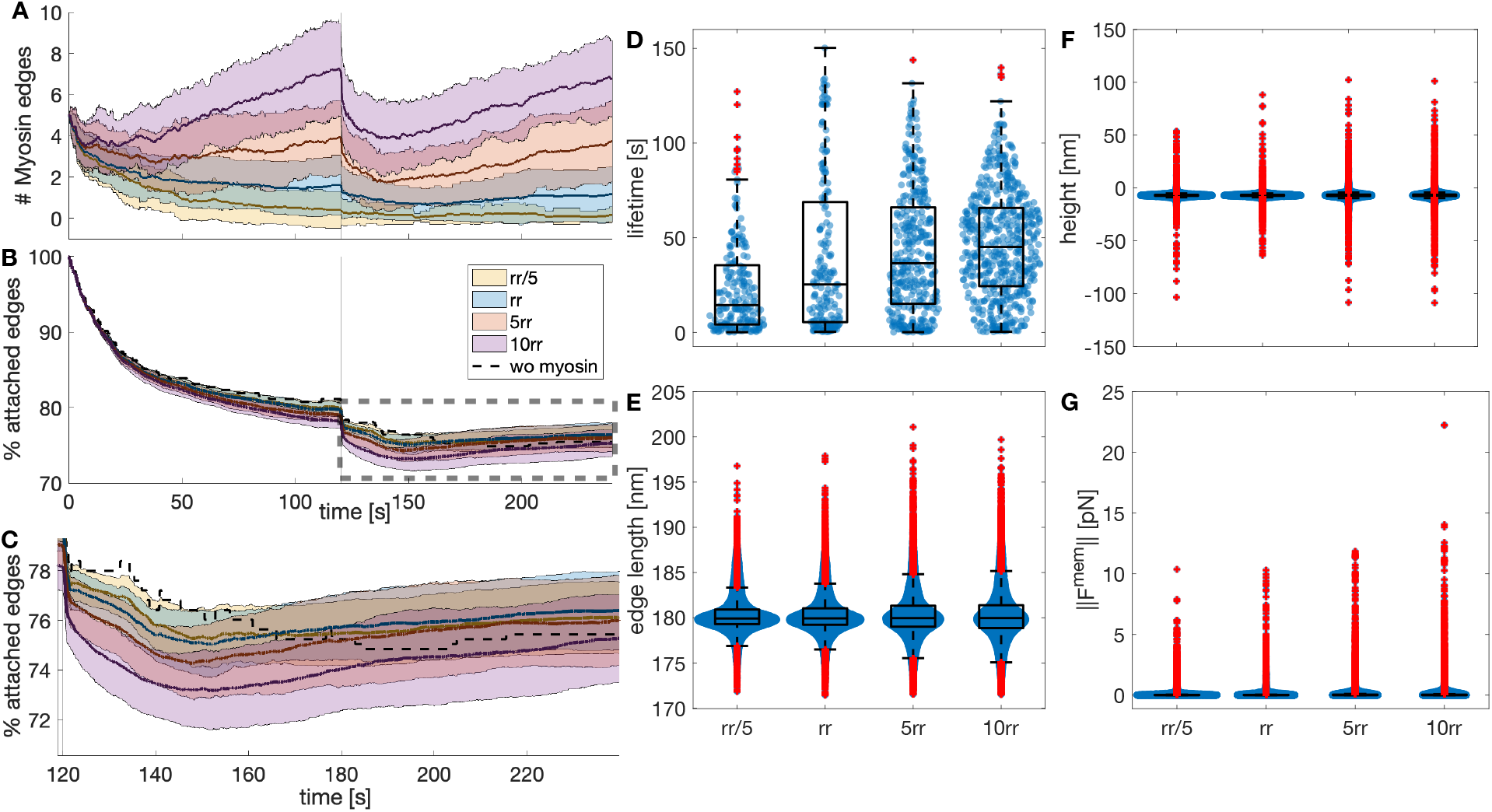
Myosin dynamics under different stochastic addition and removal rates. **A**) Temporal evolution of the myosin edges in the actin-spectrin meshwork with different ratio of addition and removal rates (*rr* = *φ*_*a*_*/φ*_*b*_), color-coded as in B. The values correspond to *rr* = *φ*_*a*_*/*(5*φ*_*b*_), *φ*_*a*_*/φ*_*b*_, 5*φ*_*a*_*/φ*_*b*_, 10*φ*_*a*_*/φ*_*b*_. The thick line represents the mean and the shadowed area is the standard deviation from 31 simulations. **B**) Temporal evolution of the percentage of attached spectrin edges over time. **C**) Zoom image of the rectangle in B. **D**) Boxplot of myosin edges lifetime. Boxplot of spectrin edges length (**E**), F-actin nodes height (**F**), and magnitude of the force generated by the membrane (**G**) at the end of the simulation for different *rr*.

We examined the final configuration of the simulations. We found that the spectrin edges length (Fig. 6E) and actin node height (Fig. 6F) were less spread for the original parameters than for the increased *rr*. For the actin node height, the interquartile range (IQR) was 0.9242 for *rr*, 1.3378 for 5*rr*, and 1.3324 for 10*rr*. For the spectrin edge length, IQR = 1.8201 (*rr*), 2.3228 (5*rr*), and 2.5261 (10*rr*). With larger *rr*, the spread of the magnitude of the membrane force also increased from an IQR = 0.0158 to IQR = 0.0398 for 5*rr* and IQR = 0.0339 for 10*rr*. Therefore, we concluded that our original parameters, which resulted in a single myosin per 2.5 μm^2^ acting on the meshwork after releasing it from focal adhesions, gave a more efficient recovery. More or fewer myosin edges in the meshwork did not improve the recovery of spectrin edges and produced stress in the meshwork, i.e., the spectrin edge length deviates more from the resting length and the height of the actin node is more divergent, which exerts spring and bending energy. These qualitative findings suggest that cells use the required number of myosins to enhance cytoskeletal recovery after inducing stress and this function may be tightly regulated.

### 2.5 Cell adhesion promotes actin-spectrin meshwork stabilization and conserves cell shape

Cells in suspension and adhered cells have different mechanical properties (*37*). Therefore, we next investigated how the actin-spectrin meshwork differs in cell-like geometries in suspension, like red blood cells, and cells with adhesions, such as fibroblasts. To do this, we implemented the model on a fully connected meshwork. For cells in suspension, we chose a sphere to capture the simplest fully connected 3D shape and avoid computational challenges associated with high curvatures. In this case, unlike the meshwork resembling a patch of membrane, the initial configuration of the spectrin edges was under stress due to deviations in their resting length (Fig. 7A). Such deviations were necessary to obtain a spherical shape. However, during the first few seconds of the simulation, the edges with smaller lengths than the resting length were removed (blue line, Fig. 7D,E). After 360 seconds of the simulation, we observed that the sphere crumbled (Fig. 7A) and increased its membrane force (Fig. 7B), while dynamically adding and removing spectrin edges (blue line, Fig. 7E,F). Moreover, the sphere volume (blue line, Fig. 7C) and the number of myosin edges (blue line, Fig. 7F) showed a sustained decrease, arising from the myosin contractile action. However, experimental data show that myosin contractility maintains cell shape (*18*). Thus, we hypothesized that there must be further mechanisms that guarantee cell shape maintenance.

**Figure 7:**
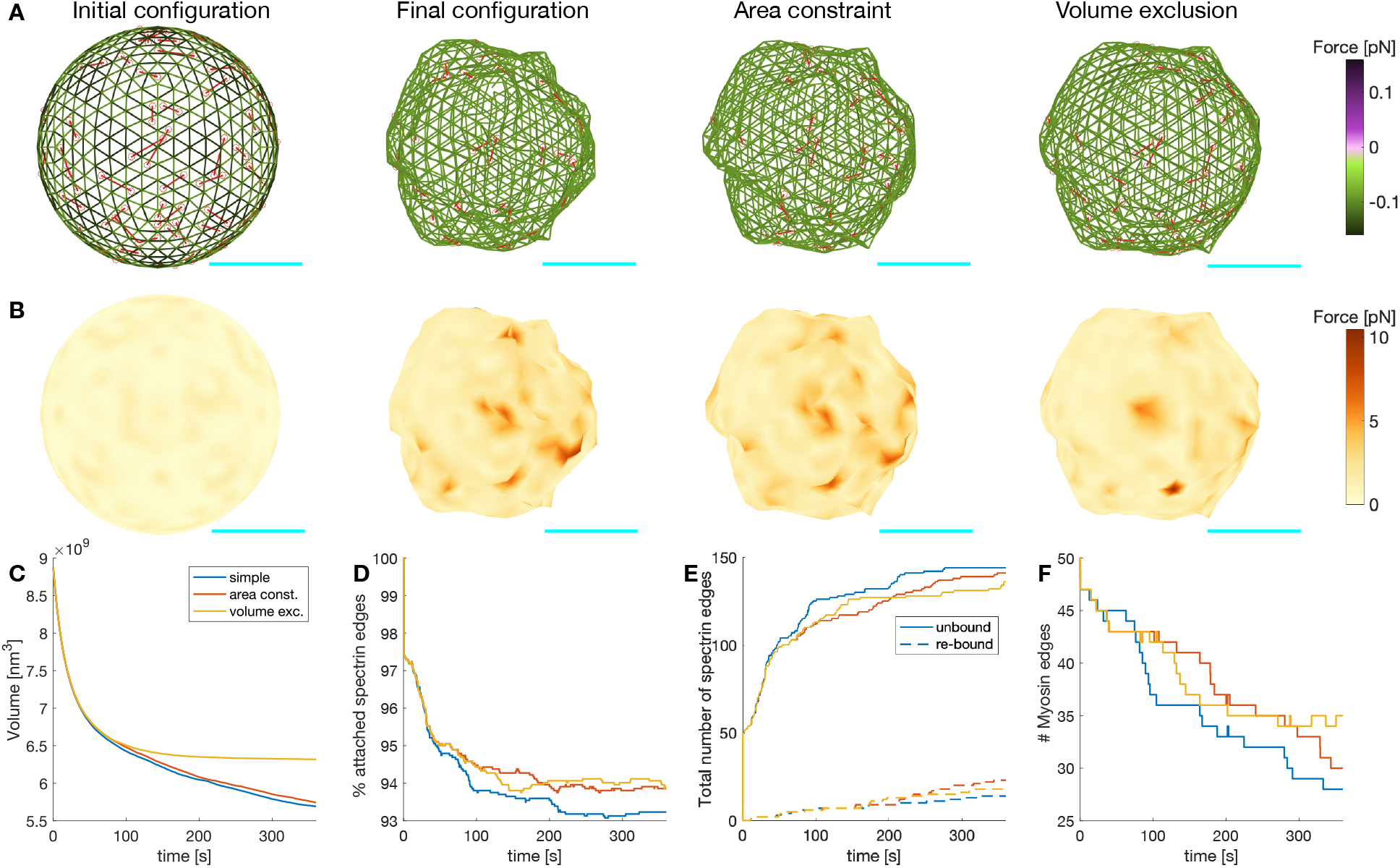
Actin-spectrin meshwork dynamics on a suspended cell. **A**) Initial configuration of the actin-spectrin spherical meshwork and 360 seconds of the simulation, with area constraint and volume exclusion. Edges are color-coded for the force generated by the spring potential energy of the spectrin edges. Red lines correspond to myosin edges. Cyan line is a scale bar corresponding to 1 μm. **B**) Meshwork in A but color-coded for the force generated by the membrane. Here, 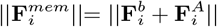. Time evolution of the volume (**C**), percentage of attached spectrin edges (**D**), total number of spectrin edges unbound and re-bound (**E**), and number of myosin edges (**F**).

As in (*38*), we assumed that the plasma membrane resists stretching. Indeed, experiments show that high stretching moduli are conserved for different types of lipids bilayers (*39*), and hence, local membrane incompressibility can be assumed (*38*). This was implemented in the model by adding a surface area constraint to the force generated by the membrane, now 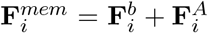 (see Module 3). We found that when the surface area was constrained, the number of spectrin edges unbound was reduced and rebinding of these edges was promoted (Fig. 7E). Thus, the percentage of spectrin edges attached was higher (Fig. 7D). We also observed that the number of myosin edges was higher (Fig. 7F), which resulted from the change in the force balance that hindered the myosin contraction, and thereby, their removal. The locations where the force generated by the membrane was high before implementing surface area constraint to the force balance, smoothed out, thereby reducing the crumbled appearance.

The size of the sphere under surface area constraint was bigger but its volume kept decreasing (Fig. 7C). It is known that nondividing adult cells maintain their size (*40*) and, based on experimental data, we only expect volume fluctuations in the absence of any stimulus at the simulation timescale (*41*). Hence, we implemented volume exclusion in the model to represent the presence of organelles in the cytosol by restricting the movement of actin nodes to a volume 15% smaller than the initial volume (Module 3). Based on experiments where hyperosmotic shocks caused a nonreversal volume decrease (*41*), we assumed that larger volume deviations trigger further cellular processes. A simulation with surface area constraint and volume exclusion showed that the sphere settled to a steady volume (Fig. 7C) while experiencing spectrin edge unbinding and rebinding events (Fig. 7E). Moreover, the force generated by the membrane was reduced (Fig. 7B). Therefore, we concluded that the interaction between the actin-spectrin meshwork and the membrane contained by surface area and volume exclusion promoted shape integrity. While the actin-spectrin meshwork allows cells in suspension to deform and bear different stresses, the membrane surface area constraint and volume exclusion guarantee shape integrity. This complements the accepted function of the spectrin skeleton in giving mechanical support to the membrane (*5*) and hints at a feedback mechanism between the membrane and the skeleton.

Most cells are embedded in the extracellular matrix and adhere to it, which alters the actin-spectrin meshwork. Therefore, we examined the meshwork dynamics in a configuration that resembles a cell adhered to a surface. We took the initial sphere configuration of Figure 8A and set *r*^*z*^ = 0 for all the F-actin nodes in the south hemisphere, i.e., locations with *r*^*z*^ *<* 0. Then, we added spring connecting edges to attach the F-actin nodes at position (*r*^*x*^, *r*^*y*^, *r*^*z*^ = 0) to fixed nodes located at (1.1*x*, 1.1*r*^*y*^, − 100 nm). Such an arrangement guaranteed that the initial length of the linker springs (*d*_*I,C*4_) was larger than the resting length (*d*_0,*C*4_). Thus, the bottom of the hemisphere was stretched during the simulation, inducing a shape change (Fig. 8A,B). We simulated the same cases as in the sphere (Fig. 7A,B) and observed that the final configuration was less crumpled when considering area constraint and volume restriction. Moreover, the spectrin edge and membrane forces were reduced, and the volume stabilized (Fig. 8A-C). The spectrin edges experienced a rapid detachment after the start of the simulation, induced by the contraction of the connecting edges (Fig. 8E). However, the percentage of attached spectrin edges immediately stabilizes (Fig. 8D). Interestingly, the number of myosins in the meshwork of the adhered cell settled to a mean value earlier than in the suspended configuration (Fig. 8F). Altogether, we found that the actin-spectrin meshwork was more stable when connected to the substrate. We hypothesize that when cells adhere, the actin-spectrin meshwork stabilizes to organize membrane proteins (*12*).

**Figure 8:**
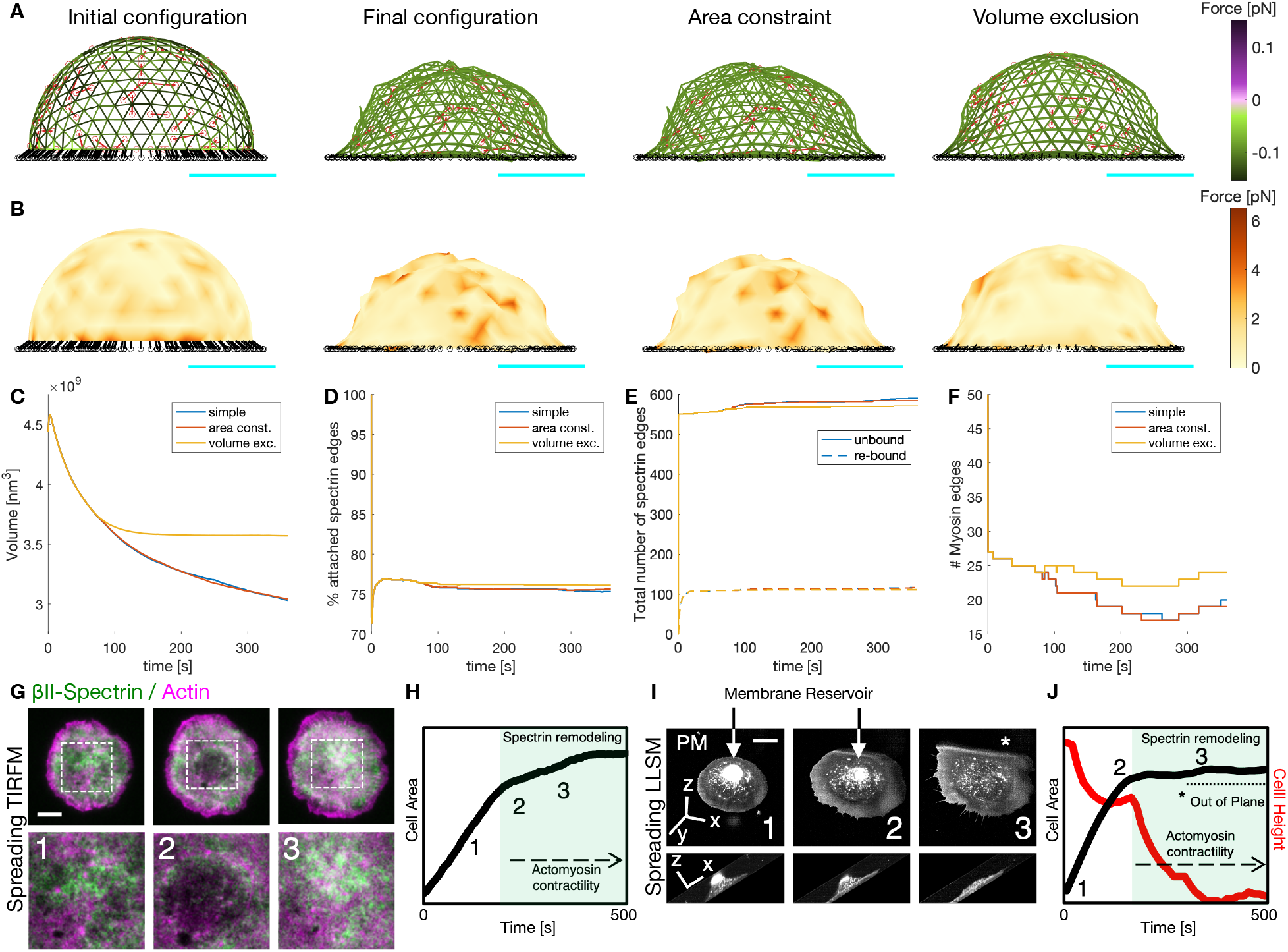
Actin-spectrin meshwork dynamics on an adhered cell. **A**) Initial configuration of the actin-spectrin spherical meshwork and 360 seconds of the simulation, with area constraint and volume exclusion. Edges are color-coded for the force generated by the spring potential energy of the spectrin edges. Red lines correspond to myosin edges. The black lines represent connecting edges and black circles, connecting nodes. Cyan line is a scale bar corresponding to 1 μm. **B**) Meshwork in A but color-coded for the force generated by the membrane. Here, 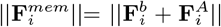. Time evolution of the volume (**C**), percentage of attached spectrin edges (**D**), total number of spectrin edges unbound and re-bound (**E**), and number of myosin edges (**F**). **G**) Cell spreading analysis at the cell body (zooms corresponding to the dashed white boxes), displayed by live TIRFM images (green: GFP-βII-spectrin, magenta: RFP-actin, scale bar: 10 μm). Relevant events observed between independent experiments are shown (1-3), in particular, endogenous actin node formation and correspondent βII-spectrin behavior. **H**) Projected Cell Area analysis over time and the relative positioning of frames 1-3 presented in G are shown in the graph. Activation of actomyosin contractility and spectrin remodeling during the slow-growth phase of spreading (P2) is highlighted in green. Figures adapted from (*10*). **I**) Cell spreading imaged by Lattice Light Sheet Microscopy in MEF transfected with the membrane reporter Scarlet-PM(Lck), scale bar: 10 μm. Relevant frames 1-3 are reported in the orthogonal view (whole cell) and in the lateral projection (to highlight cell height). The membrane reservoir is present on the top of the cell body and dissolved during the slow-growth phase of spreading (P2). **J**) Projected Cell Area (black) and Cell height (red) analysis over time and the relative positioning of frames 1-3 presented in C are shown in the graph. Activation of actomyosin contractility and spectrin remodeling during the slow-growth phase of spreading (P2) is highlighted in green, correlating to the flattening of the cell body. The portion of the cell that is excluded from the illumination plane is indicated by the asterisk (*).

Cell spreading represents an active biological process where adhesion to the substrate, membrane remodeling, and cytoskeletal modifications simultaneously occur and interplay (*42*). More specifically, previously published Total Internal Reflection Microscopy data (*10*) and novel observations by high temporal-resolution Lattice Light Sheet Microscopy (LLSM), highlighted how spectrin remodeling is driven by the re-awakening of acto myosin contractility (Fig. 8G-H). Interestingly, this slow-growth phase of spreading (also referred to as P2) corresponded to the exhaustion of the membrane reservoir (area constraint) and the flattening of the cell body towards an equilibrium state (volume constraint) highlighted by the 4D LLSM imaging approach (Fig. 8I-J). These correlative observations closely resemble the series of events captured by our model, suggesting that the enhanced stability of the adherent meshwork is important for cell function.

## 3 Discussion

Using a model of the spectrin skeleton, we examined possible mechanisms for cells to bear different stresses. Although spectrin models have been proposed (*24–26*), such a dynamic interaction between spectrin, myosin, short F-actin, and the membrane has not been previously studied. Our simulations revealed the following outcomes, relevant to the biophysics of the actin-spectrin meshwork. First, the plasma membrane is critical in lowering fluctuations in the actin-spectrin meshwork, hinting at an interaction between the plasma membrane and actin-spectrin meshwork rather than the experimentally studied function of spectrin skeleton in maintaining the stability and structure of the plasma membrane (*12*). We tested possible mechanisms that promote actin-spectrin meshwork response to different stresses and the meshwork recovery after the stresses are removed, such as spectrin unbinding and rebinding and myosin stochastic dynamics. These mechanisms are difficult to examine in experiments *in vivo* due to technical restrictions. Finally, to test the generality of our work on a membrane patch, we modeled suspended and adhered cells, which have different mechanical properties (*37*). Furthermore, we showed how these cells can conserve their shape despite the continuous turnover of spectrin and stochastic dynamics of myosin, which are necessary for responding to imposed stresses. We related our *in silico* findings with our publish (*10*) and unpublished data, where the cell size is maintained after depletion of membrane reservoir and flattening of the cell body despite the spectrin re-modeling driven by myosin contractility.

Specifically, we showed that bending energy from the plasma membrane and spectrin detachment is necessary to bear isotropic compression and shear stress. We assumed that the bending energy from the plasma membrane is transmitted to the actin-spectrin meshwork via the short-actin nodes, as in (*24*). Although more sophisticated descriptions for the link between the skeleton and lipid membrane have been proposed (*26, 28*), our chosen description of the bending energy reduces the fluctuations in the *z* plane of a simulated meshwork patch. Experimental evidence for dissociation of spectrin tetramers into dimers under shear response, which can relate to spectrin edge unbinding, has been known for a long time (*7*). However, only recently, the changes in the number of bound spectrin to short F-actin complexes have been examined using a theoretical model (*11, 31, 43*). In this work, we improved the 2D model in (*11*) by considering the rebinding of the unbound spectrin edges to test whether the system can return to the initial state after the stress is removed. Due to a lack of experimental evidence for the rebinding mechanism of spectrin bundles, we chose the simplest rule: edges rebind to the same actin nodes when the distance between the nodes is equal to the resting distance. Other rules have been proposed, for example, a model of the RBC with the stochastic addition and removal of spectrin edges shows that repeated deformations will lead to structural changes in the cytoskeleton (*43*). Future theoretical efforts can explore different rules for spectrin rebinding and the effects on the connectivity of the actin-spectrin meshwork. Furthermore, buckling of spectrin edges can be considered as on a recent model of a network of fibrin fibers, which shows the importance of buckling for describing shear response int he network (*44*).

In our simulations, the skeleton with membrane bending energy and unbinding and rebinding of spectrin edges settles to a clustered steady state when the spectrin edges recover their resting length. However, we hypothesized that, after removing the stress, the spectrin meshwork connectivity should recover. Based on the interaction between myosin and the actin-spectrin meshwork observed in fibroblast (*11*) and RBC (*18*), we added myosin to the network. As in (*11*), we assumed that myosin edges are contractile until they reach a minimum length and are removed from the network. Moreover, myosin edges are stochastically added and removed, mimicking the spatially heterogeneous contribution of myosin to cell contractility. These assumptions resulted in a more dynamic actin-spectrin meshwork, which showed an enhanced recovery from the stress. Interestingly, we found an optimum balance between the spectrin edges stochastic addition and removal rates. Experiments could test whether increasing or decreasing the number of myosin rods acting on the actin-spectrin meshwork enhances its response after some stress is induced. It has been shown that RBC contains ≈ 150 non-muscular myosin IIA bipolar filaments per cell (*18, 38*). Although a previous model of the RBC cytoskeleton considers myosin forces (*38*), it uses a deterministic description to inform the stable configurations. In our model, myosin gives a stochastic feature that allows a continuous reconfiguration of the actin-spectrin meshwork.

Next, we showed that despite the continuous stochastic dynamics of the skeleton, a fully connected meshwork can reach a stable state with a given volume and fluctuating number of spectrin and myosin edges. To keep the generality of our approach, we tested two cases that resemble cells with different properties: suspended and adherent cells. For this, we considered surface area conservation and volume exclusion, due to the presence of different molecules and organelles. Moreover, the actin-spectrin meshwork stabilizes sooner in the adhered case. Future efforts can consider how the distribution of spectrin and myosin edges changes after different stresses are applied in different cells.

In our model, we assumed that the meshwork is dynamic even when it is not stressed, in line with experimental evidence (*13*). To simulate such a dynamic meshwork, we chose a simplified representation of spectrin bundles as Hookean springs. Spectrin bundles are usually represented using a Worm Like Chain (WLC) model (*25, 28, 45*) to account for thermal fluctuations of polymers (*46*), or interpolation of the WLC (*24, 26, 47–50*) proposed by (*51, 52*), which avoids the collapse of the spectrin bundle under compression and bounds it under expansion while behaving like an ideal spring at the minimum (*47*). These representations of spectrin bundles are highly non-linear and require significant computational power. Alternatively, the simple Hookean spring potential has been used and proved (*53*) to coincide with the WLC potential used in (*47, 48*) for small extensions. In our simulations, we controlled the applied stress, which resulted in the extension and contraction of the spectrin edges within the small extension criteria (2*d*_0,*S*_ and 0.6*d*_0,*S*_, respectively) (*53*). Moreover, the resulting spectrin edges are below the spectrin length when all the repeats are unfolded ( ≈ 1022 nm) (*8*). Thus, a Hookean spring representation of the spectrin bundles is well-suited for our investigation. This mesoscopic depiction of the membrane skeleton, which omits its molecular details given in other models (*25, 30, 31*), allows us to examine the overall configuration changes due to the induced stresses. Importantly, we chose this mesoscopic model because we are interested in the effects of dynamically adding and removing, either randomly or due to applied forces, the skeleton components embedded in a membrane. The predictions derived from our model can be tested experimentally. For example, the optimum number of myosins acting on the actin-spectrin meshwork to promote its recovery after the imposed stresses are removed and the enhanced stability of the adherent cell in comparison with the suspended cell. Future efforts can add more molecular detail to our model.

## 4 Methods

### 4.1 Simulations

For the simulations in Figure 3, we solved Eq. (13) with 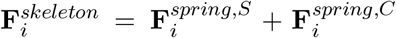 and 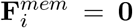 for the isotropic extension and compression and 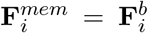 (Eq. 4) when adding bending. The unbinding and rebinding of spectrin edges were implemented first in Figure 4. Myosin dynamics are introduced in Figure 5. Hence, in Eq. (13), 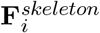 is now defined as in Eq. (10). For the plots showing the F-actin node height and membrane force, we obtain the values for each F-actin node and use interpolated coloring for the triangular surfaces.

To calculate the spreading of the data contained in the boxplot of Figure 6, we used the interquartile range (IQR), which is defined as the difference between the 75th and 25th percentiles of the data (i.e., the top and the bottom edges of the box). The IQR does not account for the data outliers and gives a better representation of the data range.

For the fully connected sphere meshwork in and semi-sphere in Figures 7 and 8, we take 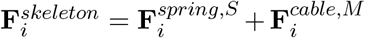 in Eq. 13. Note that when including the area constraint and volume restriction, we define 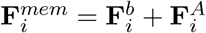.

### 4.2 Numerical Implementation

We run the simulations in MATLAB R2021a desktop computer. Following (*11*), for the patch of actin-spectrin meshwork (Figs. 3-6), we obtain the initial spectrin mesh with the MATLAB’s `delaunayTriangulation.m` function and implement the forward Euler method to solve Eq. (13). We trace the sphere in Fig. 7 with the `icosphere.m` function (*54*) and use the `remeshing.m` function (*55*) to obtain a (semi)isotropic consisting of equilateral triangles with side length *d*_0,*S*_.

### 4.3 Code availability

The code will be uploaded to a public repository at the time of final publication. It will be made available to the reviewers upon request.

### 4.4 Lattice Light Sheet Microscopy

The LLSM (*56*) utilized was developed by E. Betzig and operated/maintained in the Advanced Imaging Center at the Howard Hughes Medical Institute Janelia Research Campus (Ashburn, VA); 488, 560, or 642 nm diode lasers (MPB Communications) were operated between 40 and 60 mW initial power, with 20–50% acousto-optic tunable filter transmittance. The microscope was equipped with a Special Optics 0.65 NA/3.75 mm water dipping lens, excitation objective, and a Nikon CFI Apo LWD 25 × 1.1 NA water dipping collection objective, which used a 500mm focal length tube lens. Live cells were imaged in a 37°C-heated, water-coupled bath in FluoroBrite medium (Thermo Scientific) with 0–5% FBS and Pen/Strep. MEFs were transfected 24 h before the experiment with the mScarlet-PM (Lck) plasmid (Addgene: 98821). Before the experiment, cells were trypsinized, centrifuged for 5 min at 300 × g, washed once with PBS, and serum-starved in suspension for 30 min at 37°C in CO2-independent 1 × Ringer’s solution. Suspended cells were thereafter kept at room temperature for up to 3 h. Transfected MEFs were added directly to the coverslip submerged in the media bath prior to acquisition. The time-lapse started after a positively double-transfected cell engaged with the fibronectin-coated coverslip. Images were acquired with a Hamamatsu Orca Flash 4.0 V2 sCMOS camera in custom-written LabView Software. Post-image deskewing and deconvolution were performed using HHMI Janelia custom software and 10 iterations of the Richardson-Lucy algorithm.

## Acknowledgements

LLSM imaging was performed at the Advanced Imaging Center (AIC) – Howard Hughes Medical Institute (HHMI) Janelia Research Campus. We thank John M. Heddleston and Teng-Leong Chew of the AIC for helpful discussion. The AIC is jointly funded by the Gordon and Betty Moore Foundation and the Howard Hughes Medical Institute. This work was supported by the NIH Grant Number 1RF1DA055668-01, NIH RO1GM132106, and by an Air Force Office of Scientific Research Grant FA9550-18-1-0051 to P.R., an Italian Association for Cancer Research (AIRC) Investigator Grant (IG) 207101 to N.C.G., by H2020-MSCA individual fellowship (796547) and Fondazione Cariplo Young Investigator Grant (2021-1507) to A.G.. We thank members of the Rangamani Lab for fruitful discussions.

